# The male and female gonad transcriptome of the edible sea urchin, Paracentrotus lividus: identification of sex-related and lipid biosynthesis genes

**DOI:** 10.1101/2021.08.30.458199

**Authors:** André M. Machado, Sergio Fernández-Boo, Manuel Nande, Rui Pinto, Benjamin Costas, L. Filipe C. Castro

## Abstract

*Paracentrotus lividus* is the most abundant, distributed and desirable echinoid species in Europe. Although, economically important, this species has scarce genomic resources available. Here, we produced and comprehensively characterized the male and female gonad transcriptome of *P. lividus*. The *P. lividus* transcriptome assembly has 53,865 transcripts, an N50 transcript length of 1,842 bp and an estimated gene completeness of 97.4% and 95.6% in Eukaryota and Metazoa BUSCO databases, respectively. Differential gene expression analyses yielded a total of 3371 and 3351 up regulated genes in *P. lividus* male and female gonad tissues, respectively. Additionally, we analysed and validated a catalogue of pivotal transcripts involved in sexual development and determination (206 transcripts) as well as in biosynthesis and storage of lipids (119 transcripts) in male and female specimens. This study provides a valuable transcriptomic resource and will contribute for the future conservation of the species as well as the exploitation in aquaculture settings.

**Highlights:** Assembly of a reference transcriptome of *Paracentrotus lividus* gonads.

Differential gene expression between males and female gonads of *Paracentrotus lividus.*

Identification and validation of pivotal genes involved in biosynthesis and storage of lipids.

## Introduction

*Paracentrotus lividus*, commonly known as the purple sea urchin, is the most abundant echinoid species in Europe, with an overall distribution from the Mediterranean Sea to the eastern Atlantic coast, from Scotland to Southern Morocco, Canary and Madeira Islands (Ghisaura et al. 2016). In the last few years, *P. lividus* populations have constantly decreased due to intense harvesting, habitat destruction and climate change (Bertocci et al. 2012, Bertocci et al. 2018). The significant pressure on this species, results from the high price market of their gonads, which has led to the populational collapse in some geographic areas (Fernández-Boán et al. 2012, Ouréns et al. 2015). Additional negative factors linked to populational decline, include environmental changes such as temperature, red tides and ocean acidification (Yeruham et al. 2015, Mos et al. 2016, Ohgaki et al. 2019). Economically, the purple sea urchin is highly desirable in gourmet markets, not only because their gonads are considered a delicacy but also due to their high protein and polyunsaturated fatty acid (PUFA) content, especially in omega-3 and omega-6 (Prato et al. 2018, Baião et al. 2019, Rocha et al. 2019). Additionally, the market price of sea urchin gonads is dependent on taste, colour and firmness. Testis from males have usually a sweet and milky flavour, while female ovaries taste bitter and sour (Phillips et al. 2009, Phillips et al. 2010). These characteristics together with the gonadosomatic index of sea urchins stablish the market price of the captures. The reproductive period of *P. lividus* is variable, and several factors such as diet, availability of food resources, photoperiod and water temperature can condition their spawn (Ghisaura et al. 2016). Usually, in the Atlantic coast *P. lividus* spawn is annual (Garmendia et al. 2010, Fernández-Boo et al. 2018), while in the Mediterranean Sea is bi-annual (Sellem and Guillou 2007). The reproductive apparatus of sea urchin is composed by five gonads with different colour patterns between sexes, while males present a yellow-orange pattern, female gonads are red-orange. The gonads are composed by two main types of cells: germinal cells where the gametes are produced and stored, and somatic cells defined as nutritive phagocytes which are main storage of nutrients and energetic reserves of the animal (Garmendia et al. 2010, Ghisaura et al. 2016).

Sea urchin aquaculture is based on the enhancement of gonad yield, also known as bulking. This approach consists in the capture of wild animals with the aim of improving their gonadosomatic index, gonad sensory attributes or manipulation of the reproductive cycle to commercialize sea urchin when wild animals are not available (Walker et al. 2015). Despite multiple attempts to develop intensive aquaculture practices, the poor knowledge of dietary composition for juveniles and the period of time necessary to reach market size are the main bottlenecks in *P. lividus* production (James et al. 2015, Liu and Chang 2015). Moreover, sea urchin aquaculture has been hampered by the lack of genomic resources, with some minor examples involving the production of sea urchin triploids to increase the roe size by suppression of gametogenesis and the enhancement of nutritive phagocyte growth (Böttger et al. 2011, Walker et al. 2015). Importantly, to select parental individuals with valuable genetic traits (e.g. gonad size, colour, or lipid content), for production purposes, it is crucial to develop *omic* resources of the species.

Decisively, in last few years several strategies have been applied to study and comprehend multiple traits with value in production context (Liu et al. 2005, Wang, Ding et al. 2020). Next generation sequencing approaches, both at genomic and transcriptomic level, are now established to explore general biological species features, whether from the commercial or fundamental point of view (Sea Urchin Genome Sequencing 2006). For instance, transcriptome studies in different sea urchin species have been conducted recently to investigate the genes involved in the synthesis of polyunsaturated fatty acids in *Strongylocentrotus nudus* (Jia et al. 2017, Wei et al. 2019), the sex related genes in *Mesocentrotus nudus* (Sun et al. 2019) or the response to ocean acidification in *S. purpuratus* (Evans et al. 2017). Other species with less economic value, but highly important to local communities, have been also studied and information regarding genes involved in development, fertilization, toxin effects or immune system response against pathogens are now available (Gaitan-Espitia et al. 2016, Laruson et al. 2018). In line with transcriptomic studies, the number of whole genome sequencing projects in sea urchin species has drastically increased after the massive effort in the genome sequencing of *S. purpuratus* (Sea Urchin Genome Sequencing 2006, Janies et al. 2016, Kinjo et al. 2018, Davidson et al. 2020). In *P. lividus*, genomic and transcriptomic resources are scarce in public databases. The few studies available have explored the embryonic development an experimental model for evolutionary and ecotoxicological fields (Gildor et al. 2016, Ruocco et al. 2016, Chassé et al. 2018, Tato et al. 2018, Galasso et al. 2019, Morroni et al. 2019); also, the proteins involved in the attachment by their tube feet were studied for their use in industrial and medical purposes (Pjeta et al. 2020). Regarding the gonadal tissue of adult specimens, only two articles were released, one studying the protein patterns of gonads in males and females at different stages of development (Ghisaura et al. 2016), and second measuring the gene expression level of different pollution biomarkers after metal exposure (Di Natale et al. 2019).

Here, we reported an in-depth transcriptome analysis of the gonads from three adult males and females. These analyses establish the basis for new studies on gonad development, sex differentiation, growth and also lipid and colour traits. Through functional annotation and phylogenetic analyses, multiple pivotal genes involved in sex determination and differentiation, gametogenesis, lipid metabolism and PUFA biosynthesis were scrutinized in this species. This study will be highly valuable not only to improve the information of this keystone species in the Atlantic and Mediterranean Sea, as also could be used to improve and boost the sea urchin aquaculture in Europe.

## Material and methods

### Experimental setup, RNA extraction and sequencing

Ten adult sea urchins were collected in July 2018 in Vila Chã beach (41.295160 N, 08.737073 W) by scuba diving (68.6 ± 4.9 mm diameter; 123.10 ± 22.82 g weight; 13.51 ± 2.29 % gonadal somatic index) (Fig.1a). Initially, four females and six males were identified after gonad extraction by visual identification in a light microscope. A small piece of the gonad was immediately stored in an Eppendorf with 1 ml of RNA later (Sigma) at 4°C for 24 hours and then, frozen at −80°C until RNA extraction. Other piece of the gonad was stored for histological analysis according with Rocha et al. (2019). Sea urchin histological sections were evaluated with light microscopy and all females were in gonadal stage IV and all males in gonadal stage III according to Machado et al. (2019). For RNA extraction, 50-100 mg of gonad tissue was homogenized in 1 ml of trizol (NZY Tech, PT) using a Precellys homogenizer (Bertin Inst., France). RNA was extracted according manufacturer’s instructions. RNA concentration was measured in a DeNovix DS-11 spectrophotometer (DeNovix Inc, USA) and 4 μg of RNA were subjected to DNAse treatment (RQ1 - Promega) according manufacturer’s instructions. After DNAse treatment, RNA was isolated and cleaned using the Total RNA isolation kit (NZY Tech, PT) according manufacturer’s protocol and diluted in a final volume of 40 μl in mQ water. Finally, the concentration was measured again in a DeNovix DA-11 spectrophotometer. RNA integrity was observed in a 2% agarose gel and the best 3 males and 3 females were selected for RNA-Sequencing. RNA integrity and quantity were evaluated using the Agilent 2100 Bioanalyzer (Agilent technologies, Santa Clara, CA, USA). The RNA integrity number was >6.1 in all samples. At the end, six samples (3 females and 3 males) were shipped to Novogene (Honk Kong) Company Limited, and sequenced using Illumina HiSeq-4000 platform (150×2bp, paired-end, 30 million sequencing reads).

**Figure 1.**
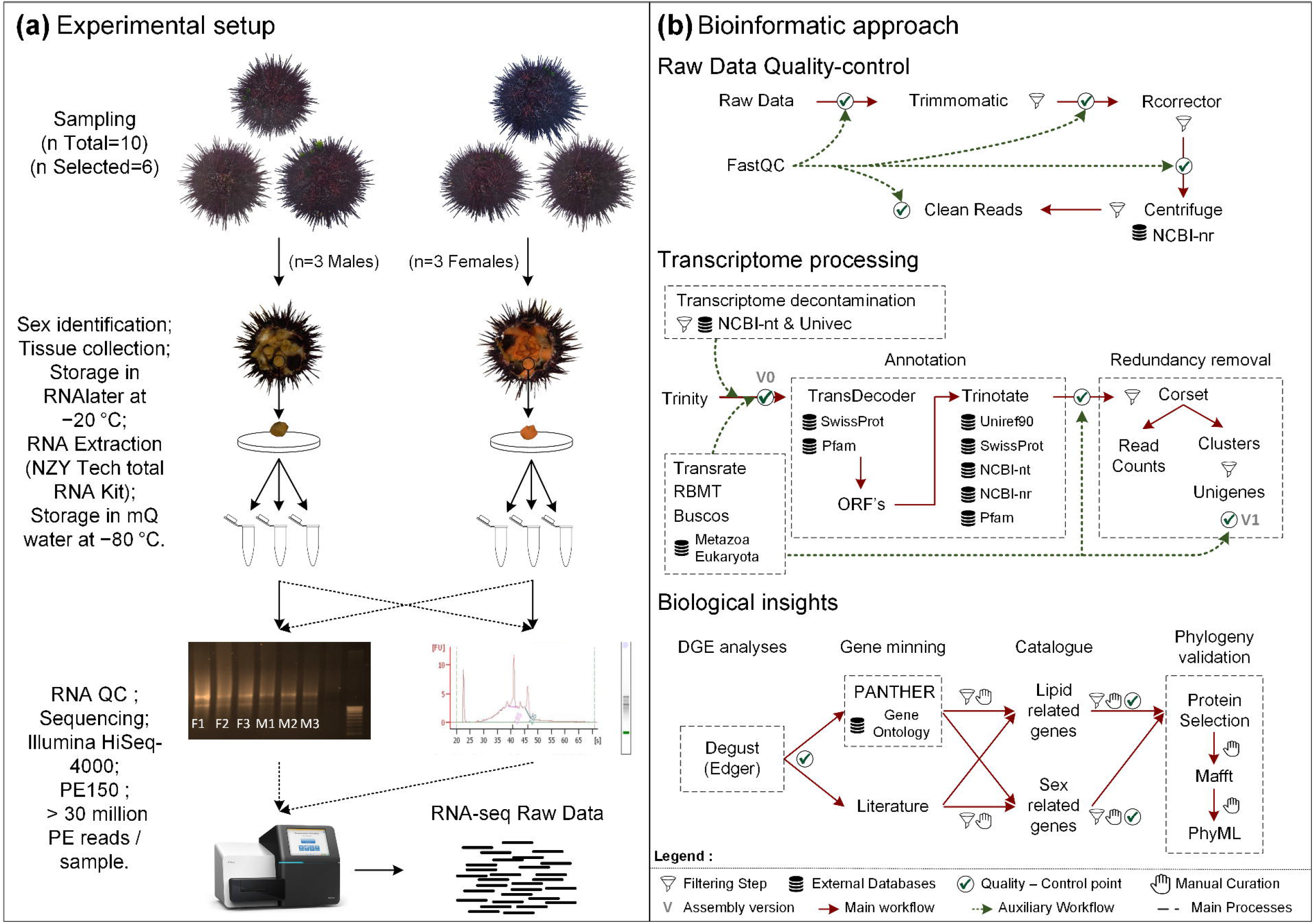
Sampling and bioinformatic workflow of the study. **(a)** Sampling and experimental setup used to perform RNA extraction and sequencing. **(b)** Bioinformatics workflow used to process the raw datasets, perform the differential gene expression and obtain biological insights.

### Raw data clean up

Initially, the RNA-Seq quality profile, of each sample, was assessed with FastQC (v.0.11.8) (http://www.bioinformatics.babraham.ac.uk/projects/fastqc/). After that, Trimmomatic (v.0.38) (Bolger et al. 2014) was used to trim and drop reads with quality scores below 5, at leading and trailing ends, with an average quality score below 20 in a 4 bp sliding window and with less than 36 bases length. Next, the error correction method, Rcorrector (v. 1.0.3) (Song and Florea 2015), was applied to correct random sequencing errors, with the default settings. In the final of the clean-up process, still it was used the Centrifuge (v. 1.0.3-beta) (Kim et al. 2016) software (Fig.1b). On this approach, all corrected raw reads were screened against the NCBI-nt database (ftp://ftp.ccb.jhu.edu/pub/infphilo/centrifuge/data/) (v.nt_2018_3_3) to obtain a taxonomic classification of each contig, with a minimum hit length of 50. Reads labelled by Centrifuge as non-Echinodermata (NCBI: taxid7586) were considered to be contaminants and excluded from the dataset.

### *De novo* assembly transcriptome, decontamination and quality assessment

After quality filtering control, clean reads were first concatenated and then assembled using Trinity (v. 2.8.4) (Grabherr et al. 2011, Haas et al. 2013) with the specific parameter (SS_lib_type RF). To remove possible sources of contaminations, all transcripts were blasted against NCBI-nt (Download; 20/11/2019) and Univec (Download; 02/04/2019) databases using Blast-n tool (v. 2.9.0) (Fig.1b). Transcripts with a match to *Echinodermata* taxon (NCBI: taxid7586), with an e-value cut-off of 1e-5, identity score of 95 % and minimum alignment length of 100bp, or without matches at all in NCBI-nt database, were retained. Transcripts matching other taxa than *Echinodermata* in NCBI-nt our Univec database were considered exogenous to the *P. lividus* transcriptome (Assembly V0) and exclude from this dataset.

### Functional annotation

The Trinotate pipeline (v. 3.0.2) (https://github.com/Trinotate/Trinotate) was used to perform the functional annotation of the transcriptome (Bryant et al. 2017). In first step, the TransDecoder software (v.5.3.0) (https://transdecoder.github.io/) was applied to predict the open reading frames (ORFs) with at least 100 amino acids. Next, both aminoacid and nucleotide sequences of Assembly V0, were blasted (blast-n tool (v. 2.9.0) (Altschul et al. 1997) and blast-x /p tools of DIAMOND software (v 0.9.24) (Buchfink et al. 2015)), against non-redundant database of NCBI (NCBI-nr) (v. 20/01/2020), NCBI-nt (Download; 30/03/2019), Swiss-Prot (Download; 18/02/2020) (The UniProt Consortium 2016), Uniref90 (Download; 04/09/2019) (Suzek et al. 2007) databases and searched in Pfam (Download; 18/02/2020) (Punta et al. 2012), eggnog (Powell et al. 2011), Kyoto Encyclopedia of Genes and Genomes pathways (Kanehisa and Goto 2000). The report generated by the Trinotate pipeline was filtered by an e-value cut-off of 1e-5 (Fig.1b).

### Transcriptome filtering and redundancy removal

The transcriptome filtering and redundancy removal were performed based on several criteria. First, the functional annotation and ORF prediction of the Assembly V0 version were scrutinized. Thus, all transcripts with at least one blast hit (blast - n, x or p in Uniref90, NCBI-nt or NCBI-nr Databases) in an *Echinodermata* species or codifying to a protein were collected (Annotated Assembly V0) (Fig.1b). Second, the raw reads were mapped onto the Annotated Assembly V0 with Bowtie2 (v. 2.3.4.2) (Settings; --no-mixed --no-discordant --end-to-end --all -- score-min L,-0.1,-0.1) and filtered with Corset (v.1.0.9) (Davidson and Oshlack 2014) software. While all the transcripts containing at least 10 reads mapping were kept, the transcripts with few reads mapping were considered spurious and discarded. In addition to the filtering by read coverage, the Corset software clustered of transcripts, based on the ratio of shared reads and expression patterns. Finally, and to remove the redundancy, only the longest transcript per cluster (Unigene) was collected (Assembly V1) (Fig.1b). To evaluate the quality of both transcriptome versions (V0 and V1), several strategies were applied. Completeness and gene content of the transcriptomes were assessed using Metazoa and Eukaryota lineage-specific profile libraries of Benchmarking Universal Single-Copy Orthologs tool (BUSCO v. 3.0.2) (Simão et al. 2015). Accuracy was assessed by the percentage of original clean sequence reads mapped against the Assembly V0 (RMBT) using Bowtie2 (v. 2.3.4.2) (Langmead and Salzberg 2012) with default settings. Structural integrity and general stats of the transcriptome were examined using TransRate (v. 1.0.3) (Lang-Unnash 1992) with default settings(Fig.1b).

### Differential gene expression analyses

Differential gene expression (DGE) analyses were performed using the Unigenes matrix of raw counts, generated by Corset software during the filtration steps, and the Degust (v.4.1.1) platform (Powell 2019) (http://degust.erc.monash.edu/). All Unigenes containing less than 1 CPM (count per million mapped reads) in at least three samples were removed from the dataset. In addition, the multidimensional scaling plot was applied to check the variance of the samples. Thereafter, the edgeR (v.3.26.8) package (Robinson et al. 2009) of R (v.3.6.1) was used to perform the DGE analyses between males and females samples of *P. lividus*. During the DGE analyses, the trimmed mean of M-values (TMM) method (Robinson and Oshlack 2010) was applied to perform the normalization of the values across the samples. Finally, the Unigenes were considered differentially expressed if the values of False Discovery Rate - corrected (FDR) p-value < 0.05 and log2|fold change| ≥ 2. The heatmaps were performed in Heatmapper Expression tool (http://heatmapper.ca/expression/), within the Average Linkage clustering method and the Euclidean Distance measurement method (Babicki, Arndt et al. 2016)

### Gene Ontology analyses and gene cataloguing

To perform the gene ontology (GO) analyses we used the blast-x annotations of the differentially expressed genes (DGEs) against the Uniref90 database. To do that, two datasets were used; firstly, the total number of DGEs; second only the upregulated genes in males and females. Technically, the Uniref90 Id’s were subjected to Panther v.15.0 for gene function classification using the sea urchin *Strongylocentrotus purpuratus* as a model organism (Mi et al. 2019). Identification of sex and lipid related genes was done by searching specific GO keywords (Sex related terms: reproductive process GO:0022414; reproduction GO:0000003; developmental process GO:0032502; signalling GO:0023052 and Lipid Related terms: biogenesis GO:0071840; metabolic process GO:0008152; catalytic activity GO:0003824) in panther output and trinotate report, as well as, different genes already described in the bibliography involved in the sex differentiation and lipid related pathways in Echinodermata phylum.

### Phylogenetic analysis

Full-length amino acid (aa) sequences were used for phylogenetic analysis, to reduce the impact of nucleotide composition bias. The lipid and sex-related genes as well full-length aa sequences were selected accordingly with their relevance within the main pathways previously identified by Panther and after an exhaustive analysis of the related literature (Suppl. Table. 1). After collecting the target sequences, orthologous proteins were gathered from NCBI database, using the blast-p algorithm with e-value cut-off of 1e-5 and at least 75% identity in echinoderms (*Strongylocentrotus purpuratus, Apostichopus japonicus, Asterias rubens*, and *Crassostrea gigas*) and vertebrates model species (*Danio rerio, Xenopus tropicalis, Gallus gallus, Mus musculus*, and *Homo sapiens*) (Suppl. Table. 1). Multiple alignments were performed with all orthologs sequences using MAFFT v7.402 free software (Katoh et al. 2019) with the L-INS-I algorithm (Katoh and Standley 2013). Subsequently, the alignments were manually reviewed, and short sequences and gaps were removed. To remove gaps we used the GapStrip / Squeeze v2.1.0 (http://www.hiv.lanl.gov) software and all columns of the multi alignment with 95% nucleotides fulfilled were kept for the phylogenetic analyses. Phylogenetic trees were constructed using PhyML 3.0 server (Guindon et al. 2010), using the maximum likelihood method with the default parameters (specific substitution model details in Suppl. Table. 2). Later the phylogenetic trees were visualized by Dendroscope (Huson et al. 2007) and rooted with mollusc sequences from *Crassostrea gigas*.

## Results and Discussion

### Histological analysis

In the case of females, the gonads presented an advanced stage of development being two of them at stage III (F1,F2) and one at stage IV (F3) according to scale of Byrne (1990). Stage III is considered a premature stage where all oocytes are at all stages of development. The nutritive phagocytes are displaced from the central position to the periphery by the large oocytes (Fig. 2 a, b, c). The stage IV is a mature stage with ovaries closely –packed and most of them with a diameter higher than 90 μm. The NP at this stage are in a thin layer around oocytes (Fig. 2 c). All males were in stage III of maturation. These stages are characterized by mature testes packed with spermatozoa (S) and nutritive phagocytes (NP) are clearly limited to the periphery (Fig. 2 d, e, f). The collection of individuals in a similar stage of maturation allowed to obtain a comparative dataset between males and females.

**Figure 2.**
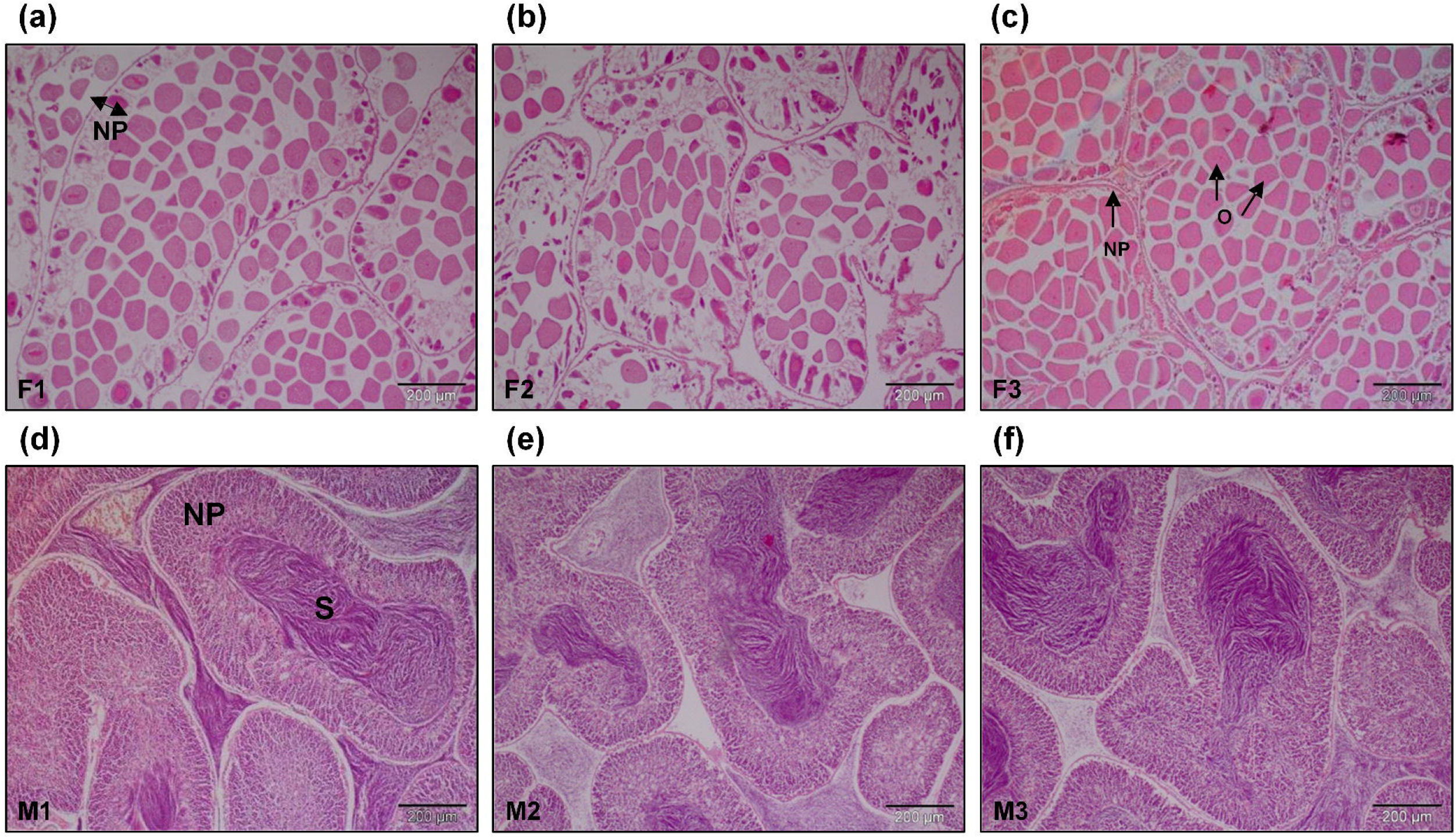
Histological sections of gonads used for transcriptome sequencing. Females were in stage III (a, b) and IV (c) of maturation. All males were in stage III of maturation. Scale bar (200 μm). NP: nutritive phagocyte; O: oocyte; S: spermatid.

### Sequencing data

From the ten *P. lividus* specimens collected, six (three males (M1, M2, M3) and three females (F1, F2, F3)) were selected to perform gonadal RNA-Seq sequencing. The RNA-Seq approach generated a total of 204,865,648 PE reads that after validation with several quality-control and filtering softwares, were reduced to 203,895,472. Although the conservative approach used to perform the clean-up, a high percentage of raw reads (99.53% of the initial datasets) showed phred score□≥□Q20 which indicates the high quality of the initial dataset. In the end of this process, all clean the raw reads were deposited to the NCBI database and can be consulted under the BioProject accession: PRJNA625933 (Table. 1a).

### The *de novo* assembly and filtering

To build the transcriptomic reference of *P. lividus*, we applied *de novo* assembler Trinity. Briefly, the six samples of *P. lividus* were pooled together and inputted in the Trinity assembler as a unique dataset. Next, the Trinity software selected about of 44,665,675 (21.91%) reads during the normalization stage and produced the first draft assembly of the transcriptome. Importantly, this version of the transcriptome assembly, was submitted to an extensive quality control against the NCBI-nt and UniVec databases, which allowed to remove sources of contaminations and to build the first version of the gonadal transcriptome of *P. lividus* (Assembly V0) (Table. 1b). Unexpectedly, this transcriptome version transcriptome showed a huge number of transcripts (more than 1 M), a N50 transcript length of 720bp and a total length size of 756,496,790bp. Biologically, these values can be explained by several factors. For instance, features such interspecific variation and heterozygosity can impact the building of *de novo* assembly references. On the other hand, high repeat content can confuse the *de novo* assemblers leading to high rates of fragmentation, spliced transcripts in many sub-transcripts, and production chimeric or artefactual transcripts (Lima et al. 2017). These characteristics were raised in the genome projects of the sea urchins, *Strongylocentrotus purpuratus, Hemicentrotus pulcherrimus* and *Lytechinus variegatus*, in which the authors have found high levels of heterozygosity as well as repeat content and several difficulties in the building of the genome references. To remove the redundancy and mitigate the impact of these values in further analyses, we next applied several filtration steps. These filters have used three main approaches, codifying status, functional annotation and read coverage of the transcripts.

The codifying status of the transcripts was determined with the TransDecoder software. From 1,310,553 initial transcripts, only 138,992 transcripts codify to an ORF with 100 or more aminoacids. The aminoacid and nucleotide sequences were then functional annotated using a set of different databases (Fig 3a). After searches in several databases, the Uniref90, NCBI-nt and NCBI-nr databases were selected to filter the transcriptome and about of 185,644 transcripts were found to have at least one blast hit in an echinoderm species. Thus, the Annotated version of the transcriptome (Annotated Assembly V0), was obtained though the sum of coding transcripts and transcripts with blast hit but without an ORF (225,700 transcripts - 138,992 protein coding transcripts; 86,708 additional transcripts with blast hit). Subsequently, the 225,700 transcripts were inputted in the Corset software to remove all transcripts with scarce evidences of read mapping and to cluster the remaining transcripts based on multi-mappings. In the end, Corset grouped 141,138 transcripts (>10 reads mapping) in the 53,865 clusters / Unigenes- (Assembly V1). Both versions of the transcriptome assembly as well the annotation reports and open read frames can be consulted in figshare digital repository (Link to the reviewers - https://figshare.com/s/e6d85154080ad008ee7c).

**Figure 3.**
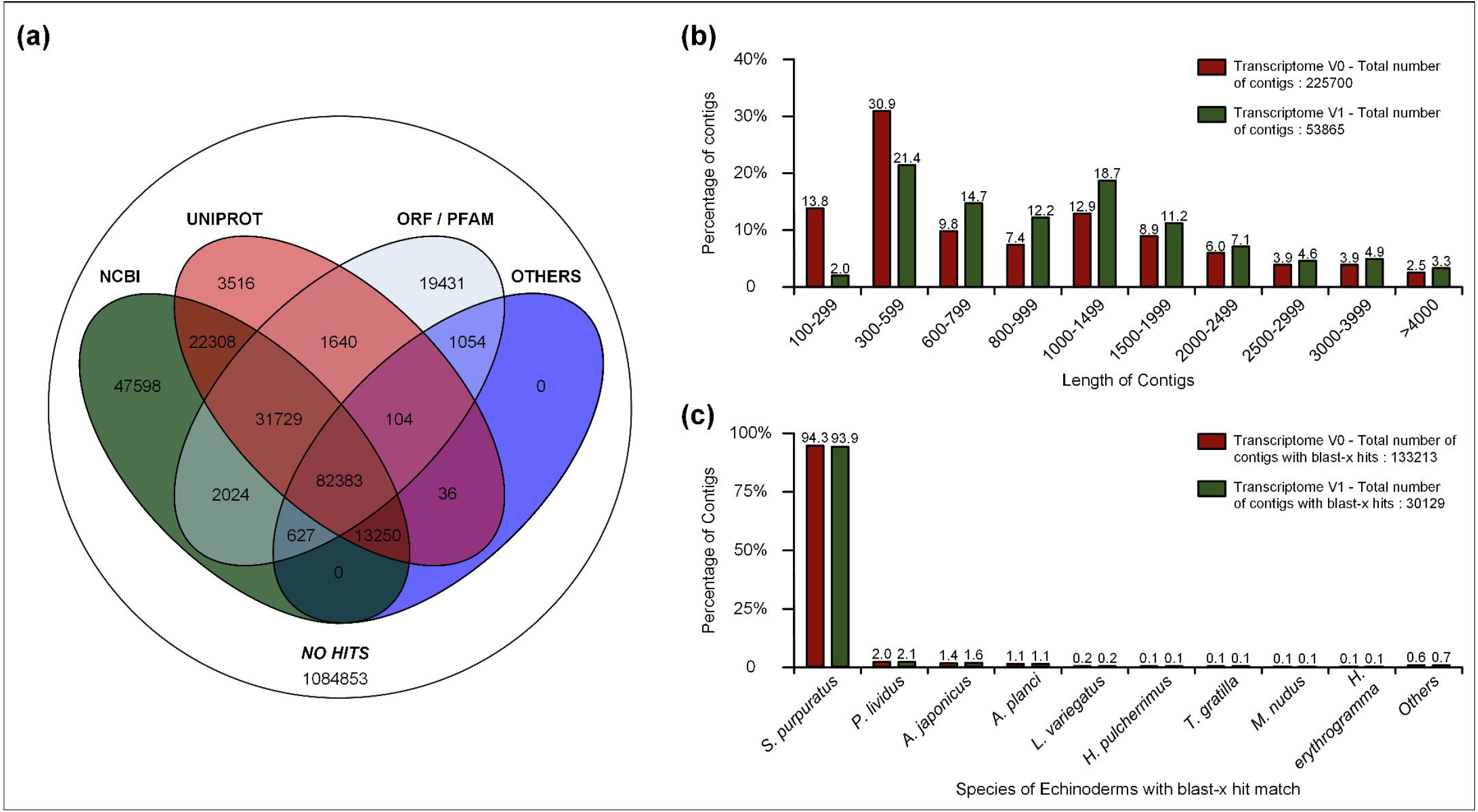
Functional annotation and clean-up analyses of the V0 and V1 versions of transcriptome assembly of P. lividus. **(a)** Venn diagram of the transcripts (V0 transcriptome) codifying to an open reading frame or matching with NCBI, UNIPROT and PFAM databases. **(b)** Length distribution of the transcripts in versions V0 and V1 of the transcriptome assembly. **(c)** Blast-x analysis of the V0 and V1 versions of the P. lividus transcriptome assembly.

Overall, this strategy has significantly reduced the size and the redundancy of the transcriptome assembly, and is similar to previous analyses with sea urchins transcriptomes (Chen et al. 2015, Gaitan-Espitia et al. 2016, Gaitán-Espitia and Hofmann 2017, Wong et al. 2019, Zhang et al. 2019, Zhang et al. 2019, Shi et al. 2020). The final version (Assembly V1) of *P. lividus* transcriptome had only 4,11% of the initial number of transcripts (53,865), an N50 transcript length of 1,842, more than the double of the first version, and 1/10 of the initial transcriptome length (Table. 1b). Importantly, both transcriptome versions (V0 and V1) were carefully inspected, assessed and compared with three methods.

First, the gene content was analysed using the BUSCO tool. Searching the two libraries profiles, Eukaryota and Metazoa, we observed a high completeness of the transcriptome assemblies. While in V0, 99.4 and 98.1 % of the total gene groups were found in both databases, in the V1 we identified 97.4 and 95.6 % respectively. Importantly, all BUSCOs analyses showed a high percentage (>92%) of genes classified as complete, and almost no gene content was lost during the filtration process. On the other hand, the number of missing genes in both versions per database is remarkably low (V0, Eukaryota - 0.6 %, Metazoa - 1.9%; V1, Eukaryota - 2.6 %, Metazoa - 4.4 %). Comparing these values with others in literature, we can conclude that *P. lividus* gonadal transcriptome is one of the most complete in sea urchins reported to date (Table.1b). For instance, in the sea urchins *Strongylocentrotus intermedius* (Zhan et al. 2019), in *Echinometra* sp. embryos (Uthicke, Deshpande et al. 2019) or in giant red sea urchin *Mesocentrotus franciscanus* (Gaitán-Espitia and Hofmann 2017), *de novo* assemblies achieved 82.5%, 91%, 88% of complete genes in eukaryotic database, respectively.

Second, we performed a structural comparison between V1 and V0 assemblies. The Fig 3b showed a clear decrease in the percentage of short contigs (< 600 bp) from the V0 (43.7%) to the V1 (22.4%) of the transcriptome. Consistently, in the remaining classes the V1 version showed always higher percentages of transcripts than V0 version, indicating that small contigs were largely affected by the filtration steps, while the percentages of the large contigs remaining almost unmodified.

In the final test, the rate of back mapping reads to V0 and V1 transcriptomes was analysed. As expected, the RBMT decrease 10,31% from the V0 to the V1 (Table.1b). Using the structural and RBMT analyses it is possible to conclude that a part of the reads not mapped in the V1 version belongs to small and fragmented contigs removed during the filtration set. This results is coherent with the biological features of sea urchins, that with high repetitive regions lead to high percentages of fragmented and spliced transcripts in several sub-transcripts. Lastly, the top 10 Echinodermata species with highest number of transcripts matching the NCBI-nr database in both V0 and V1 Assembly versions are shown in Fig.3c. As can be consulted, both versions displayed a very similar distribution.

### Functional annotation

The functional annotation was performed to the V0 version of the gonadal transcriptome and thereafter gathered to the V1 version. Overall, the V0 version showed only 15.75 % (206,543) of the total transcripts annotated against at least one database, and 6.29 % annotated in all databases (Fig.3a; Table.1b). On the other hand, the V1 dataset of Unigenes presented a ratio 91.05% (49,048) of annotated transcripts (Table. 1b, Suppl. Table. 3). Once the remaining analyses explored the V1 assembly. The following results of functional annotation will be focus on this version. In KEGG, eggNOG, and Pfam databases we found between 33.17 and 39.61 % of the Unigenes. On the other hand, in the general databases of NCBI-nt, NCBI-nr, UNIREF90 and Swiss-Prot were found 31,257; 34,584; 35,692 and 22,939 matches. The blast-x results against the NCBI-nr database showed 34,537 match hits in 615 species. Of these, 30,129 (87.24%) Unigenes matched against species of Echinodermata, 597 against Cnidaria, 189 against Mollusca, 184 against Arthropoda and 3061 against others Phyla. Among the top 10 Echinodermata species with highest number of unigenes, the *S. purpuratus* species stand out the remaining species, with 93.9 % of the hits (Fig.3c). Although several genomes and transcriptomes of sea urchins are available (e.g. Sea Urchin Genome Sequencing 2006, Cameron et al. 2015, Cary et al. 2018, Kinjo et al. 2018, Davidson et al. 2020), to the date of this analyses only *S. purpuratus* species has a uniformized genome annotation, CDS and proteins available in NCBI databases. In addition, the *P. lividus* and *S. purpuratus* are close related species in terms of phylogeny, which makes this result expected (Mongiardino Koch et al. 2018). The remaining eight species of Fig.3c have between 2 and 0.1 % of the hits and comprise one sea star, *Acanthaster planci* (Class: Asteroidea), one sea cucumber, *Apostichopus japonicus* (Class: Holothuroidea) and seven other sea urchins (Class: Echinoidea).

### Differential expression analyses

The differential gene expression analyses between the male and female samples were performed in the Degust platform. After submitting the matrix raw of counts to the platform, all Unigenes were filtered by the CPM (1 CPM in at least 3 samples). This filter allowed gathering only Unigenes with strong read support (17,581). Next, it was evaluated the variance across the samples using the multidimensional plot. Importantly, both female and male samples are clearly grouped in two independent clusters. Moreover, the female samples presented higher variance than male samples (Suppl. Fig. 1). The DGE analyses yielded a total of 6722 Unigenes with p-value < 0.05 and log2|fold change| ≥ 2. Overall, were found 3371 Unigenes up regulated in *P. lividus* males and 3351 up regulated in *P. lividus* females (Suppl. Fig. 2, Suppl. Table. 4). Moreover, 163 and 162 Unigenes were found to be female and male specific, respectively (Suppl. Table. 5, 6). Regarding the annotation status of the DGE Unigenes (DGEs), about 5278 DGEs codifying to an ORF, and 6267 have a match hit annotation in UNIPROT or NCBI databases.

### Gene ontology analyses

The ontology analyses included all DEG and were mainly focused on the Biological Process category. These analyses showed most of the genes related with cellular process (35%), metabolic process (23.8%) and biological regulation (14.9%), while only a few directly involved in reproduction (0.8%) or reproductive process (0.8%) (Suppl. Fig.3). When the gene ontology annotation was determined per sex, once again most of the genes were related with cellular process (32.6% F vs 37.3% M), metabolic process (24% F vs 23.7% M) and biological regulation (15.8% F vs 14% M), while reproduction has once again a lower expression (0.3% F vs 1.2% M) (Suppl. Fig.4). Remarkably, the reproduction gene ontology term has 4 times more expressed genes in males than females. In terms of developmental process occurs the opposite, with more DGE present in females (4% F vs 1.8% M) (Suppl. Table 7; Suppl. Fig.4).

Next, to identify candidate genes involved in sex differentiation, determination and gonad development of *P. lividus*, the DGE genes were filtered using key GO terms and literature in the Echinodermata phylum. According to Panther classification, only 32 entries were associated to reproduction (GO:0000003) and reproductive process (GO:0022414) gene ontology terms. When the Trinotate report was scrutinized, a higher number of genes linked to gonad development and sex differentiation (GO:0008584, GO:0007548) were founded. The Spermatogenesis category (GO:0007283) was the most abundant category with 115 entries all of them founded up-regulated in males, corresponding to the 57.21% of the total sex related genes founded in males (Suppl. Table 8). The second category with more entries was Spermatid development (GO:0007286) with 2 up-regulated genes in females and 49 in males. Then, oogenesis (GO:0048477) appeared with 7 entries in females and 10 in males. After that, males gonad development (GO:0008584) with 13 entries 6 in females and 7 in males; fertilization (GO:0009566) with 11 entries all of them in males; oocyte maturation (GO:0001556), 9 entries, 1 in females and 8 in males; males meiosis (GO:0007140), 5 entries, 1 in females and 4 in males; gonad development (GO:0008406), 4 entries, 2 in females and 2 in males; females gonad development (GO:0008585), 3 entries all in males and finally oocyte development (GO:0048599) with 2 entries, one in females and another in males. The highest number of entries were related with lipid metabolic process (GO:0006629) with 30 entries, 15 in females and 15 in males; then, lipid transport (GO:0006869) with 21 hits, 14 in females and 7 in males was the second most abundant group; cholesterol metabolic process (GO:0008203) with 13 hits, 7 in females and 6 in males; lipid glycosylation (GO:0030259) with 12 hits all in females and lipid storage (GO:0019915) with 10 hits, 5 in females and 5 in males were the most abundant groups. Additionally, some genes were involved in fatty acid elongation (GO:0030497) and fatty acid elongase process (GO:0009922) were found (Suppl. Table 9).

Initially, the total number of genes found in female and male gonads was similar. However, after filtering a higher number of genes related with sex and reproduction was detected in males (Fig.4a). Moreover, the DGE present in males was 3.28 times higher than in females (154 vs 47). In contrast, in lipid related genes a total of 119 genes were obtained, 48 hits of genes up-regulated in males and 68 in females. In this case, as expected, the highest number of genes involved in fatty acid production was founded in sea urchin females (Fig.4b).

**Figure 4.**
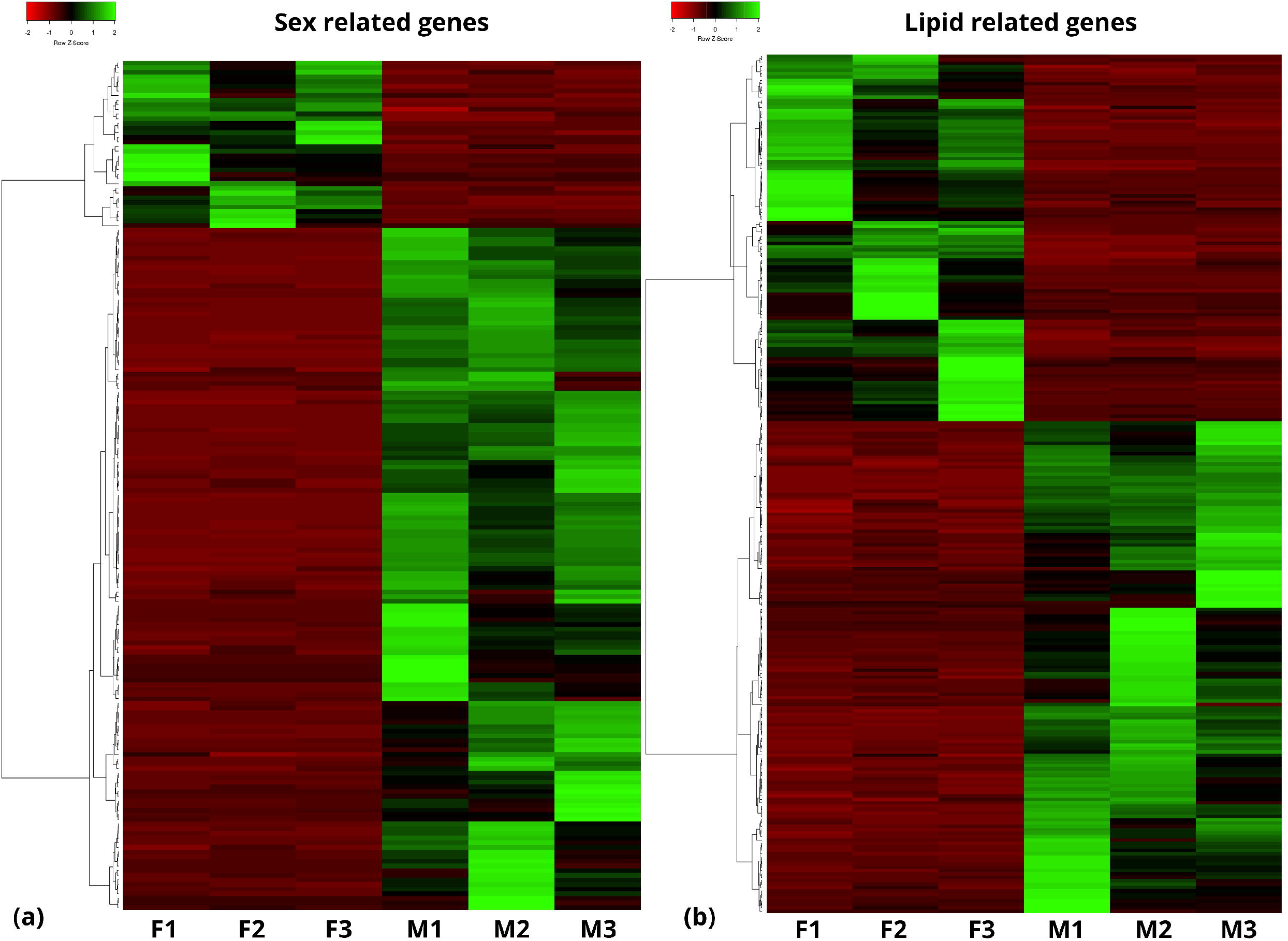
Heatmap of up and down regulated genes. **(a)** Sexual development and determination genes. **(b)** Lipid related with biosynthesis and storage of lipids.

### Catalogue Sex and Lipid-related Unigenes

#### Unigenes involved in sexual development/determination

As previously described, sea urchins are diploid species with heteromorphic chromosome sex mechanisms of the XY genes (Lipani et al. 1996). A characteristic of *P. lividus* is the lower number of chromosomes (2n=36) in comparison with other echinoid species (2n = 42 to 44) (Lipani et al. 1996). Usually, sex determination in gonochoric species such as *P. lividus* involves a robust transcriptional regulation (Nef et al. 2005). Interestingly, in the majority of these species the male specimens have a higher number of genes being expressed in the gonads (Nef et al. 2005, Tao et al. 2013, Teaniniuraitemoana et al. 2014, Shen et al. 2020). Notwithstanding, there are some species where the opposite also occurs (González-Castellano et al. 2019, Piprek et al. 2019).

Sex differentiation includes several processes regulated by a high variety of genes and transcription factors. Among these genes, Sox genes are some of the well described in the literature as key factors for sexual determination. In *P. lividus* we found five Sox genes differentially expressed (Suppl. Table 8). While Sox30 (DN1680_c0_g1_i5) was only found up-regulated in males, other Sox genes such as SoxB1, Sox21 and Sox4 (DN7136_c0_g1_i2; DN7136_c0_g2_i4; DN18728_c0_g1_i1) were uniquely found up-regulated in female gonads. Sox30 also known as Sex determination factor Y has been described as a key regulator of transcription in mouse spermatogenesis (Zhang et al. 2018), suggesting an important role in *P. lividus* testis development. On the other hand, the transcription factor SoxB1, one of the most expressed genes in *P. lividus* female gonads, is described as predominantly expressed in females and can have a function of oocyte protection and regulating the expression of ZP proteins (Yue et al. 2015). Despite, the finding of several Sox genes, other important genes such as Sox9, a major transcription factor for testis development (Zhang et al. 2018), was not found in *P. lividus*. Although unexpected, this result is coherent with the literature, where this and other important genes did not show differential gene expression in testis or ovaries of sea urchins’ species (Sun et al. 2019). Moreover, in Echinodea class seems to exists different Sox genes being differential expressed in gonads. Interestingly, in *Mesocentrotus nudus* species four different Sox genes such as Sox1, 3, 6 and 9 were found (Sun et al. 2019) and none of them was founded in *P. lividus* transcriptome. These differences can be explained by the differential developmental stages of the gonads in the different studies (Yue et al. 2015).

The Dmrt1 gene (DN11344_c0_g1_i5), usually involved in testis development, was also found up-regulated in *P. lividus* testis. As in many other species, including sea urchins, Dmrt1 is described as playing a major role in sex determination and differentiation in several vertebrate and invertebrate species (Zhang and Zarkower 2017, Nagasawa et al. 2019, Sun et al. 2019). In accordance with the report of (Sun et al. 2019) in *M. nudus* species, also the Nanos (DN9673_c0_g1_i2) gene in *P. lividus* is up-regulated in female gonads. Although Nanos has a key role in embryonic development in sea urchin (Fujii et al. 2006), in adult ovaries has a pivotal role in proliferation and survival of germline stem cells and cyst development (Forbes and Lehmann 1998, De Keuckelaere et al. 2018). Additionally, several genes related with the Notch and Delta pathway (e.g Neurogenic locus notch homolog protein 1 and 2 (DN2510_c0_g1_i1; DN17254_c0_g1_i4); notch gene (DN9493_c1_g2_i3), notch ligand (DN44109_c0_g2_i3) and Delta gene (DN44109_c0_g2_i3) were found in female gonads. While in adults this pathway is an essential regulator of cell proliferation during development and oogenesis (Feng et al. 2014, Irles et al. 2016, Sun et al. 2019, Zhang et al. 2019), in sea urchin embryos is related with the development process regulation (Materna and Davidson 2012). Another gene involved in the developing gonads and granulosa cells in adult mice is the GATA-type Zinc finger protein 1 (GLP-1) (Li et al. 2007, Pangas and Rajkovic 2015). This gene, not characterized in *S. purpuratus*, also was expressed in *P. lividus* females (DN31305_c0_g1_i2) and previously described in *M. nudus* (Sun et al. 2019). Importantly, this gene seems to have a crucial role in the regulation of oocyte meiosis and formation of primordial follicles, once the GLP-1 knock-out in both males and females mice results in infertility (Li et al. 2007).

Regarding the spermatogenesis regulation, several genes such as DMC1 (DN19900_c0_g1_i5), Spo11 (DN82908_c0_g1_i1), Hsd3 (DN1732_c1_g1_i2) and two SPATA genes, 24 and 45 (DN632_c1_g1_i2; DN847_c0_g1_i3), were found up-regulated in male gonads. DMC1 is a recombinase essential for meiosis in male testis, regulating the homologous recombination of chromosomes. Depletion of DMC1 leads to malformation of sperm, or aneuploidy spermatocytes in several species (Chen et al. 2016). DMC1/LIM15 (DN19900_c0_g1_i5) from *S. purpuratus* were founded in testis of *P. lividus* indicating that as in other eukaryotes, the expression of DMC1 is essential for the development of the spermatocytes in sea urchin. Also, Spo11 is essential in meiotic recombination by generating double strand breaks and it is required for meiotic synapsis (Romanienko and Camerini-Otero 2000). Two genes of SPATA family were found in the transcriptome, SPATA24 and SPATA45. The spermatogenesis-associated (SPATA) family consists in several genes with critical roles in spermatogenesis. For instance, the knock-out of these genes in humans led to several problems in sperm motility, sperm production and germ cell development (Sujit et al. 2020). Finally, several testis-specific serine/threonine protein kinases (Tssk) (e.g tssk1, 4 and 5; DN1880_c1_g1_i12; DN20093_c0_g1_i3; DN2835_c2_g1_i6) were detected in the gonadal transcriptome, mainly associated to the male gonad. In general, these genes are associated with the spermatogenesis regulation and they are widespread across the eukaryota domain, including sea urchins (Wang et al. 2016, Sun et al. 2019).

#### Lipid Unigenes determination

The main components during gametogenesis in *P. lividus* female gonads are proteins, but lipids also play a remarkable role in the viability of gametes (Sanna et al. 2017). Lipids, such as phospholipids (PL) and cholesterol, are structural components of cell membranes and the key to somatic growth (Liu et al. 2007). It increases the synthesis and storage of lipids during gonadal development, reaching maximum values in stage IV and in males in stage III (final stage), mainly for energy reservoirs (Sanna et al. 2017).

Several genes involved in the biosynthesis of long-chain polyunsaturated fatty acid (LC-PUFA) during gametogenesis, such as putative fatty acid elongation protein 3 (up-regulated in males) (DN12877_c0_g1_i2), elongation of very long chain fatty acids protein 6 (DN6887_c0_g2_i2), or fatty acyl desaturases A (Fads A) (DN230613_c0_g1_i7), were identified in the *P. lividus* transcriptome. Fads A mediates the introduction of unsaturation (double bond) into a fatty acyl chain (Guillou et al. 2010), showing Δ5-desaturated activity in which it uses 20: 3n-6 and 20: 4n-3 to transform into ARA and EPA (Kabeya et al. 2017). LC-PUFA biosynthesis activity was also reported in other sea urchin’s species such as *Strongylocentrotus intermedius* (Han et al. 2019). The gene expression of Fads A in the gonad is congruent with ARA and the EPA as the most abundant LC-PUFAs in the *P. lividus* gonad (Sanna et al. 2017). Different acyl-CoA ligase family members involved in long-chain-fatty-acid-CoA ligase (LC-PUFA-acid CoA ligase) such as LC-PUFA CoA ligase 1 and 6 (DN12809_c0_g1_i8 and DN79126_c1_g1_i1 respectively) were up-regulated in gonads of females while the LC-PUFA CoA ligase 3, 4 and 5 (DN3246_c0_g1_i1, DN3764_c0_g1_i13, and DN12660_c0_g1_i4) were found up-regulated in gonads of males (Suppl. Table. 9).

Other gene involved in the LC-PUFA incorporation into the mitochondria, mainly through the hydrolysis of triglycerides (García-Rincón et al. 2016), was carnitine O-palmitoyltransferase 2 (DN25985_c0_g2_i1). This Unigene have showed higher expression in female gonads than in males of *P. lividus*. Consistently, this gene showed crucial activity in transcriptome analysis for fatty acid metabolism of other sea urchins such as *Strongylocentrotus intermedius* (Wang et al. 2019).

Furthermore, PUFAs play a fundamental role as the main components of complex molecules such as phospholipids or triglycerides. An example of intermediate formation products is found in the sn1-specific diacylglycerol lipase beta that catalyzes the hydrolysis of arachidonic acid (AA) esterified diacylglycerol (DAG) to produce the main endocannabinoid, 2-arachidonoylglycerol (2-AG), which can be further cleaved by downstream enzymes to release arachidonic acid (AA) for cyclooxygenase (COX) mediated eicosanoid production. Also, DAG is an intermediate in the glycerolipid and glycerophospholipids metabolism for the biosynthesis of TAG and PL. In male gonads, high expression of sn1-specific diacylglycerol lipase beta (DN17156_c0_g2_i1) was showed.

Phospholipids are part of the cell membrane and essential for cell growth (Byrd 1975). Most of the phospholipids are incorporated through the diet, but *P. lividus* has the capacity for its endogenous biosynthesis (Byrd 1975). Thus, phosphatidylcholine (PC) is one of the major phospholipids in the sea urchin (Mita et al. 1994). Importantly, the Kennedy pathway route is used for phospholipids (Gibellini and Smith 2010). Cholinephosphotransferase 1 (chpt1) is the last enzyme in charge of obtaining PC from 1,2-diacyl-sn-glycerol and CDP-choline (EC 2.7.8.2). Differential expression analyses showed how cholinephosphotransferase 1 (DN72675_c0_g2_i1) was up-regulated in female gonads as an active pathway in the last PC biosynthesis step. In contrats, a significantly lower chpt1 activity was identified in oocytes of *Arbacia punctulata* compared to sperm (Ewing 1973). Furthermore, this study results showed a higher expression of lysophosphatidylcholine acyltransferase 2 (DN14424_c0_g1_i10) in male gonads, as an alternative route for the biosynthesis of PC as a final product. The enzyme encoded by lysophosphatidylcholine acyltransferase 2 plays a role in phospholipid metabolism, specifically in the conversion of lysophosphatidylcholine to phosphatidylcholine in the presence of acyl-CoA (Law et al. 2019). The alternative pathways to obtaining PC between males and females may be related to the composition of PUFA and LC-PUFA in positions sn-2 and sn-1,3. Thus, in females of *P. lividus* gonads, the sn-1,3 position of LC-PUFA such as ARA or EPA determines the use of TAGs in response to acclimatization or involvement in reproduction, respectively (Sanna et al. 2017). Also, the male gametes of the sea urchin use PC and TAG as energy (Mita et al. 1994), while in the gonad, PC is incorporated into the cell structure (Ewing 1973).

Several genes involved in TAG biosynthesis, such as diacylglycerol kinase zeta (DN39457_c0_g1_i2), nuclear envelope phosphatase-regulatory subunit 1 (DN10024_c0_g1_i6), microsomal triglyceride transfer protein large subunit (DN3378_c1_g1_i6), are presented in gonadal transcriptome. Most of the genes related to triglycerides’ metabolism were up-regulated in females gonadal concerning to the males. Thus, TAGs are the highest lipid class in gonads in *P. lividus*, varying seasonally due to lipid turnover after each period (Sanna et al. 2017). Similarly, during the egg development of the sea urchin, *Anthocidaris crassispinu*, TAGs are stored and later used in the early stages of development (YASUMASU, HINO et al. 1984). In contrast, in spermatozoa, the TAGs proportion is low or maybe absent (Mita et al. 1994). This is consistent with the higher activity of triglycerides genes related to gonadal development in *P. lividus* female than male gonads.

Furthermore, several genes related to cholesterol metabolism, transport, and binding such as scavenger receptor class B member 1 (DN1029_c0_g1_i3), NPC intracellular cholesterol transporter 1 (DN7961_c0_g1_i6), low-density lipoprotein (LDL) receptor-related protein 6 (DN74960_c0_g2_i3), and Apolipoprotein D (DN8923_c0_g2) were identified (Suppl. Table 9). A higher expression of these genes was found in female gonads compared with males, with related functions as a membrane receptor, transport, and binding of high-density lipoprotein cholesterol (HDL), low-density lipoprotein (LDL), and cholesterol. Moreover, genes such as sterol O-acyltransferase 1 have increased activity in male gonads. This enzyme plays a role in lipoprotein assembly and dietary cholesterol absorption (Yang et al. 1997). Thus, in female ovary and *P. lividus* spermatozoa, cholesterol was identified as a relevant lipid component (Mita et al. 1994, López-Hernández et al. 1999).

### Phylogenetic validation

One of the biggest challenges in gene ortholog identification is related to alterations in gene repertoire among species, as a consequence of loss, duplication, or the appearance of paralogs genes (Altenhoff and Dessimoz 2009). Here, we phylogenetically validated 12 relevant target genes annotated in our transcriptome, with theirs orthologs full-length aa sequences of other species of echinoderms sea urchin (*Strongylocentrotus intermedius*), starfish (*Asterias rubens*) and, sea cucumbers (*Apostichopus japonicus*), vertebrate model species fish (*Danio rerio*), amphibians (*Xenopus tropicalis*), birds (*Gallus gallus*), mammals (*Mus musculus* and *Homo sapiens*) and mollusks (*Crassostrea gigas*). This is a powerful method to detect annotation and assembly errors caused during the transcriptome assembly for non-model species (Guang et al. 2021). Genes with functional characterization, also validated and described in the literature for echinoderms were included in the analysis such as Fads, PPARs and Gonadotropin-Releasing Hormone-Type (Kabeya et al. 2017, Tian et al. 2017, Capitão et al. 2020). Our results showed that, all target lipid and sex-related full-length aa sequences clustered into a strongly supported clade of echinoderms with the orthologs of sea urchin (*Strongylocentrotus intermedius*), starfish (*Asterias rubens*), and sea cucumbers (*Apostichopus japonicus*) (Suppl. Fig. 5). A similar topology was identified in almost all trees (Suppl. Fig. 5), one group of vertebrates, other from echinoderms, and the last one of molluscs (*Crassostrea gigas*). As expected genes involved in lipid metabolism such as Fads A and chpt1 were clustered in the echinoderm clade with 99% posterior probability support and 100% with the other sea urchin (*Strongylocentrotus intermedius*) (Suppl.Table 5). The study of Kabeya et al. (2017) performed the functional characterization of *P. lividus* Δ5 desaturase (Fads A) and validated by phylogenetic analysis using molluscs, other sea urchins (*Strongylocentrotus purpuratus* and *Lytechinus variegatus*), and other species of marine echinoderms such as lilies (*Oxycomanthus japonicus*), sea cucumbers (*Sclerodactyla briareus*) and starfish (*Patiria miniata*). Supported by the results of Kabeya et al. (2017), we were able to obtain a similar result in our phylogenetic analysis that validates the functional annotation of key target genes in our transcriptome.

## Conclusion

To conclude, the gonad transcriptome from both sexes of sea urchins lead to the identification of a larger number of genes involved in sex determination, sex differentiation, gonad development, spermatogenesis and oogenesis and also in several pathways of lipid metabolism, fatty acid elongase process, lipid storage and cholesterol metabolic process. The results obtained in this work will provide useful data on *P. lividus* sea urchin gonad tissue and it will contribute in the future in the conservation of the species and the exploitaion for aquaculture purposes.

## Acknowledgements

This work was supported by the project CRAGIAMP – PTDC/BIA-BQM/30232/2017, co-financed by COMPETE 2020, Portugal 2020 and the European Union through the ERDF. This research was also supported by national funds “through FCT – Foundation for Science and Technology” within the scope of UIDB/0443/2020 and UIDP/04423/2020.

## Author Contributions

AMM: Data curation, Formal analysis, Investigation, Methodology, Visualization and Writing—original draft. SF-B: Data curation, Formal analysis, Investigation, Methodology, Visualization, Resources and Writing—original draft. MN: Formal analysis, Investigation, Methodology, Visualization and Writing - review & editing. RP: Investigation, Methodology, Visualization and Writing - review & editing. BC: Methodology, Validation and Writing—review & editing. LFCC: Conceptualization, Methodology, Validation, Supervision, Project administration, Resources and Writing—review & editing.

---

**Supplementary Figure 1**. Multidimensional scaling analyses. *P. lividus* male samples: M1, M2, M3; *P. lividus* male samples: F1, F2, F3.

**Supplementary Figure 2**. Heatmap of all up and down regulated genes.

**Supplementary Figure 3**. GO terms present in all DEG of *P. lividus* gonads according Panther v.15 using *Strongylocentrotus purpuratus* as model organism.

**Supplementary Figure 4**. GO terms present in all DEG of *P. lividus* gonads discriminating by males and females according Panther v.15 using *Strongylocentrotus purpuratus* as model organism.

**Supplementary Figure 5**. Phylogenetic trees were constructed by selected amino acids (aa) sequences for lipid metabolism obtained from Illumina-sequenced transcriptomes of sea-urchin gonads (*Paracentrotus lividus*) and each orthologs genes of nine invertebrate and vertebrate species. The trees were constructed using the Maximum-Likelihood Phylogenies (PhyML 3.0), the numbers at each node represent the bootstrap values as percentage and it were rooted in the *Crassostrea gigas*. Trees of sex related orthologs genes were showed as: **(a)**, fizzy-related protein homolog; **(b)**, amidophosphoribosyltransferase; **(c)**, HORMA domain-containing protein 1; **(d)**, spermatogenesis-associated protein 7 homolog; **(e)**, TBC1 domain family member 20; and **(f)**, transforming growth factor beta receptor type 3. Lipid related trees were as **(g)**, fatty acids desaturase (Fad); **(h)**, carnitine O-palmitoyltransferase 2; **(i)**, cholinephosphotransferase 1; **(j)**, peroxisome proliferator-activated receptor alpha (PPAR-∝); **(k)**, NPC intracellular cholesterol transporter 1; **(l)**, sterol O-acyltransferase 1.

